# Co-chaperone involvement in knob biogenesis implicates host-derived chaperones in malaria virulence

**DOI:** 10.1101/2021.03.11.434953

**Authors:** Mathias Diehl, Sebastian Weber, Marek Cyrklaff, Cecilia P. Sanchez, Carlo A. Beretta, Lena Roling, Caroline S. Simon, Julien Guizetti, Matthias P. Mayer, Jude M. Przyborski

**Affiliations:** Parasitology, Centre for Infectious Diseases, Heidelberg University Hospital, Im Neuenheimer Feld 324, 69120 Heidelberg, Germany; Electron Microscopy Core Facility, Heidelberg University, Im Neuenheimer Feld 345, 69120 Heidelberg, Germany; Nikon Imaging Center at Heidelberg University, BioQuant BQ001 Im Neuenheimer Feld 267, 69120, Heidelberg, Germany; CellNetworks Math-Clinic at Heidelberg University, BioQuant BQ001 Im Neuenheimer Feld 267, 69120, Heidelberg, Germany; Biochemistry and Molecular Biology, Justus Liebig University, Heinrich-Buff-Ring 26-32, 35392 Gießen, Germany; Center for Molecular Biology of Heidelberg University (ZMBH), DKFZ-ZMBH-Alliance, Im Neuenheimer Feld 282, 69120, Heidelberg, Germany

**Keywords:** *Plasmodium falciparum*, HSP40, HSP70, knobs, KAHRP, malaria, virulence, *Pf*EMP1, co-chaperone, chaperone

## Abstract

The pathology associated with malaria infection is largely due to the ability of infected human erythrocytes to adhere to a number of receptors on endothelial cells within tissues and organs. This phenomenon is driven by the export of parasite-encoded proteins to the host cell, the exact function of many of which is still unknown. Here we inactivate the function of one of these exported proteins, PFA66, a member of the J-domain protein family. Although parasites lacking this protein were still able to grow in cell culture, we observed severe defects in normal host cell modification, including aberrant morphology of surface knobs, disrupted presentation of the cytoadherence molecule PfEMP1, and a total lack of cytoadherence, despite the presence of the knob associated protein KAHRP. Complementation assays demonstrate that an intact J-domain is required for recovery to a wild-type phenotype and suggest that PFA66 functions in concert with a HSP70 to carry out host cell modification. Strikingly, this HSP70 is likely to be of host origin.

Taken together, our data reveal a role for PFA66 in host cell modification, implicate human HSP70 as also being essential in this process, and uncover a KAHRP-independent mechanism for correct knob biogenesis. Our observations open up exciting new avenues for the development of new anti-malarials.

## Introduction

*Plasmodium falciparum* causes the most severe form of malaria in humans, *malaria tropica*, responsible for over 200 million clinical cases and 400,000 deaths *per annum*, mainly in children under the age of 5 and mostly in sub-Saharan Africa^1^. The pathology associated with malaria infection is largely due to the ability of infected human erythrocytes to adhere to a number of receptors on endothelial cells within tissues and organs^1^. This cytoadherence results in reduced blood flow in the affected areas, hypoxia and (in cerebral malaria) increased intracranial pressure^1,2^. The phenomenon of cytoadherence results from parasite-induced host cell modification in which parasite-encoded proteins are transported to and exposed at the surface of the infected host cell, where they mediate endothelial binding and antigenic variation^3–5^. In addition to these surface proteins, parasites also encode, express, and export a large number of other proteins to the infected erythrocyte^6–8^. Many of these proteins are specific to *P. falciparum*, and their function is still not well understood, partly due to limitations in reverse genetic systems^8–10^.

Within the predicted ‘exportome’ are 19 proteins belonging to the family of J-domain proteins (JDPs, also called Hsp40s), and this exported family appears to be expanded in the Laveranian clade, suggesting important functions in these parasite species^8,11^. In other systems, HSP40s act as co-chaperones for HSP70, a protein family that lies at the heart of proteostasis and other essential cellular processes^12^. Previous studies have localised several members of the exported *Pf*HSP40 family to various structures within the infected erythrocyte, including red blood cell plasma membrane, Maurer’s clefts, knobs, and J-dots^10,13,14^. Knockout studies suggest that, although some of the exported HSP40s are essential, others can be deleted without any observable phenotype, and several can be deleted, resulting in aberrant cellular morphology, cytoadherence, and rigidity of infected erythrocytes, suggesting a potential role for this protein family in host cell modification and virulence characteristics^10^ (Supplementary Table 1). One exported HSP40, PFA66 (encoded by *PF3D7_0113700*, formerly *PFA0660w*), was previously localised to J-dots, novel structures within the infected erythrocyte containing further exported parasite-encoded HSP40s, an exported parasite-encoded HSP70 (*Pf*Hsp70-x), and a number of other exported proteins^13,15,16^. An earlier medium throughput knockout study failed to generate parasites deficient in PFA66, and therefore its function in the parasite’s lifecycle remains elusive^9^.

In this study, we utilise selection-linked integration-targeted gene disruption (SLI-TGD) to generate parasites expressing a non-functional PFA66 truncation mutant and characterise the resulting parasite lines. We find that inactivation of PFA66 function leads to dramatic aberrations in host cell modification, especially in knob morphology, capacity for cytoadherance and surface exposure of the virulence factor PfEMP1. Our data suggest an important role for exported HSP40s in parasite pathogenicity. Additionally, our data strongly implicate residual human HSP70 in parasite-induced host cell modification.

## Results

### Generation of PFA66 truncation and complementation cell lines

Genetic manipulation via single crossover was performed using a selection-linked integration targeted gene disruption strategy^17^ (Figure 1A). Integration of the plasmid would lead to the production of a GFP-tagged, truncated, non-functional PFA66 protein lacking the entire C-terminal substrate binding domain (SBD), referred to as dPFA. We transfected CS2 parasites that had previously been freshly selected for binding to chondroitin-sulphate-A as this binding phenotype would be essential for later characterisation^18,19^. Plasmid integration into the *PFA0660w/PF3D7_0113700* locus was verified via PCR using primers designed to yield products upon integration of the plasmid into the genome via primers spanning the integration site (Figure 1A). Appearance of bands representing the 5’ and 3’ integration, as well as the disappearance of bands representing the endogenous *PFA0660w* locus, demonstrated specific integration of the plasmid into the genome (Figure 1B), yielding parasite line CS2Δ*PFA* (referred to as Δ*PFA*). Immunodetection using an α-GFP antibody revealed the presence of dPFA::GFP fusion in cell lysates derived from Δ*PFA* but not parental CS2 parasites (Figure 1C). Together, this data indicated successful integration of the vector into the *PFA0660w* locus, leading to the production of a truncated product d*PFA* lacking the J-domain required for function. To verify that any aberrant phenotypes observed were due to a lack of PFA66 and not second site events, we generated a complementation line that expressed a full-length, functional 3xHA (hemagglutinin)-tagged copy of PFA66 from an episome, under the control of Prom^PFA66^, referred to as Δ*PFA^[PFA::HA]^*. Expression of this complementation construct was verified by immunoblotting using α-HA antibodies (Figure 1D).

**Figure 1.**
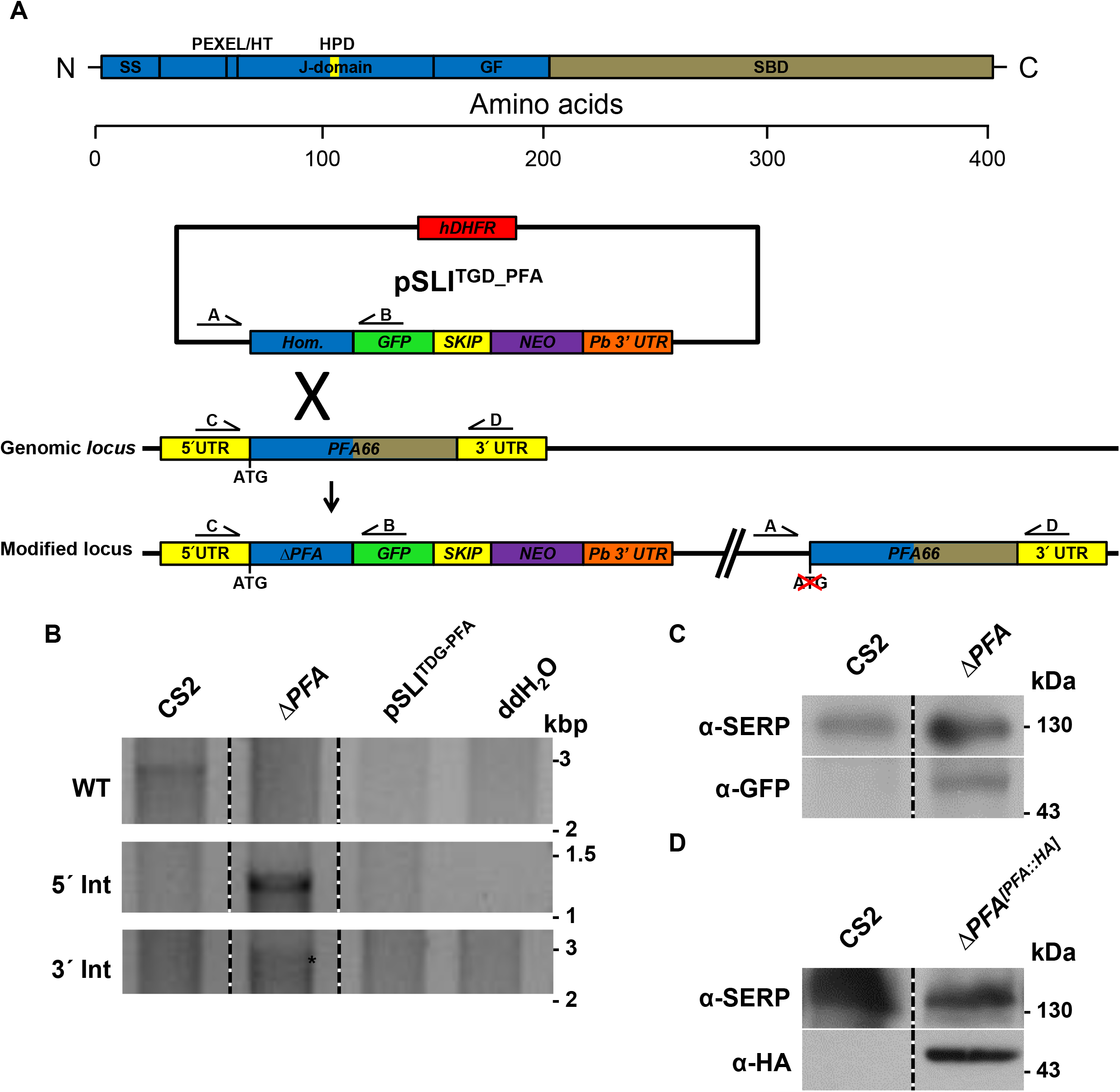
A) Strategy for inactivation of *PFA0660w* via selection-linked integration. Expression of a neomycin resistance marker (NEO) is coupled to integration of the plasmid pSLI^TGD_PFA^ into the genomic *PFA0660w* locus, leading to expression of a truncated (likely inactive) PFA66 missing its substrate binding domain (SBD). Production of PFA and NEO as separate proteins is ensured with a SKIP peptide. B) Integration PCR using gDNA extracts from the cell line Δ*PFA* and the parental cell line CS2 verifies integration of the plasmid pSLI^TGD_PFA^ into the *PFA0660w* gene in the cell line Δ*PFA*. Amplification of the wild type *PFA0660w* locus with the primers PFA0660w_5’_F and PFA0660w_5’_R is only successful in the parental strain CS2 since integration of the plasmid dramatically increases product size. PCRs using primers spanning the junctions of the integration sites (PFA0660w_5’_F and GFP_R for the 5’ region and Not-70_F and PFA0660w_5’_R for the 3’ region) demonstrate disruption of *PFA0660w*. The 3’ integration band is marked with an asterisk. Additional controls can be found in Figure S1. C) Western blot verifies truncation of *PFA0660w* in Δ*PFA*. The truncated fusion protein was detected using an α-GFP antibody, while the parasite protein SERP served as a loading control. D) Detection of a band representing HA-tagged, episomally expressed PFA66 in a western blot verifies the complementation cell line Δ*PFA^[PFA::HA]^*.

### dPFA is soluble within the host cell cytosol

Full-length PFA66 has previously been localised to the J-dots, and a follow up study suggested that the SBD of exported HSP40s is required for correct localisation^13,20^. As dPFA lacks the SBD, but still contains both an N-terminal signal peptide and a recessed PEXEL trafficking signal, we expect export of dPFA::GFP to the host cell^6,7^. Live cell imaging and immunofluorescence failed to detect dPFA, likely due to the low expression level of this protein. We therefore used equinatoxin (EQT) to selectively permeabilise the erythrocyte plasma membrane and allow sub-cellular fractionation of infected erythrocytes^21,22^. Immunodetection using antibodies against the compartment-specific markers SERP (parasitophorous vacuole), human Hsp70 (HsHSP70, red blood cell cytosol), aldolase (ALDO, parasite cytoplasm), and GFP reveals co-fractionation of dPFA with HsHSP70, verifying that dPFA::GFP is found in the host cell cytosol and furthermore that dPFA::GFP is found in the soluble phase and not in the membrane fraction as we have previously demonstrated for the full-length protein^13^ (Figure S1A). Taken together, these data suggests that deletion of the SBD of PFA66 leads to a non-functional protein.

### Truncation of PFA66 affects novel permeability pathway (NPP) activity and confers a small growth advantage

Exported parasite proteins carry out a multitude of functions supporting the survival of *P. falciparum* parasites. One of these is the establishment of NPPs of the iRBC to support the uptake of essential nutrients^23,24^. To investigate a potential role of PFA66 in NPP activity, we used a sorbitol uptake / lysis assay, which revealed that erythrocytes infected with Δ*PFA* parasites show a significantly reduced sorbitol-induced lysis, implying a reduced NPP capacity (Figure S1B). To examine whether this reduction in NPP activity affects parasite viability, growth of both cell lines was compared over four growth cycles (~8 days) using flow cytometry. Surprisingly, a slight but significant growth advantage (calculated as below 1% advantage per cycle) of the truncation cell line compared to the parental cell line was observed (Figure S1C).

### Truncation of PFA66 causes deformed knob morphology

Knobs are electron-dense protrusions of the iRBC surface that help correctly present the major virulence factor *Pf*EMP1, thus facilitating iRBC cytoadhesion and concomitantly increasing clinical pathology^3-5,25,26^. As exported HSP40s have previously been implicated in knob formation^9,14^, we used scanning electron microscopy to visualize knobs on the surface of CS2- and Δ*PFA*-infected erythrocytes. RBCs infected with the parental parasite line CS2 showed normal knob morphology, with an even distribution of small knobs over the entire surface of the infected cell (Figure 2A, S2A). Small variations in knob number and size are observed and are likely due to slightly different developmental stages of the parasite (compare 2B, C to S6A, B, D, E). In stark contrast, RBCs containing Δ*PFA* displayed, in addition to a population of normal knobs, knobs with extremely aberrant morphology. These knobs varied in their aberration, and included vastly extended knobs (Figure 2A), wide tall knobs (Figure S2), wide flat knobs (Figure S2), and branched knobs (Figure 2A). Some of the elongated knobs reached lengths of ~0.7 μm. To distinguish these abnormal structures from classical knobs, we will refer to them as *“mentula”* (plural *mentulae*). For purposes of quantification, we counted and classified knobs/*mentulae* to one of three classes I) normal/ small knobs II) abnormal/enlarged *mentulae* III) deformed/elongated *mentulae.* This analysis revealed a significant reduction in the overall number of knobs/*mentulae* in Δ*PFA*-infected RBCs when compared to wild type CS2 (CS vs Δ*PFA*, Figure 2B). 22% of surface structures exhibited abnormal morphology in Δ*PFA*-infected RBCs (Figure 2C). Interestingly, we occasionally observed extended knoblike structures on the surface of RBCs infected with CS2 parasites (Figure 2C) at a level of 2%. As a control, we complemented Δ*PFA* function with full-length PFA66 expressed from an episome, and both density and morphology of *mentulae/knobs* returned to wild type levels (Figure 2B, C).

**Figure 2.**
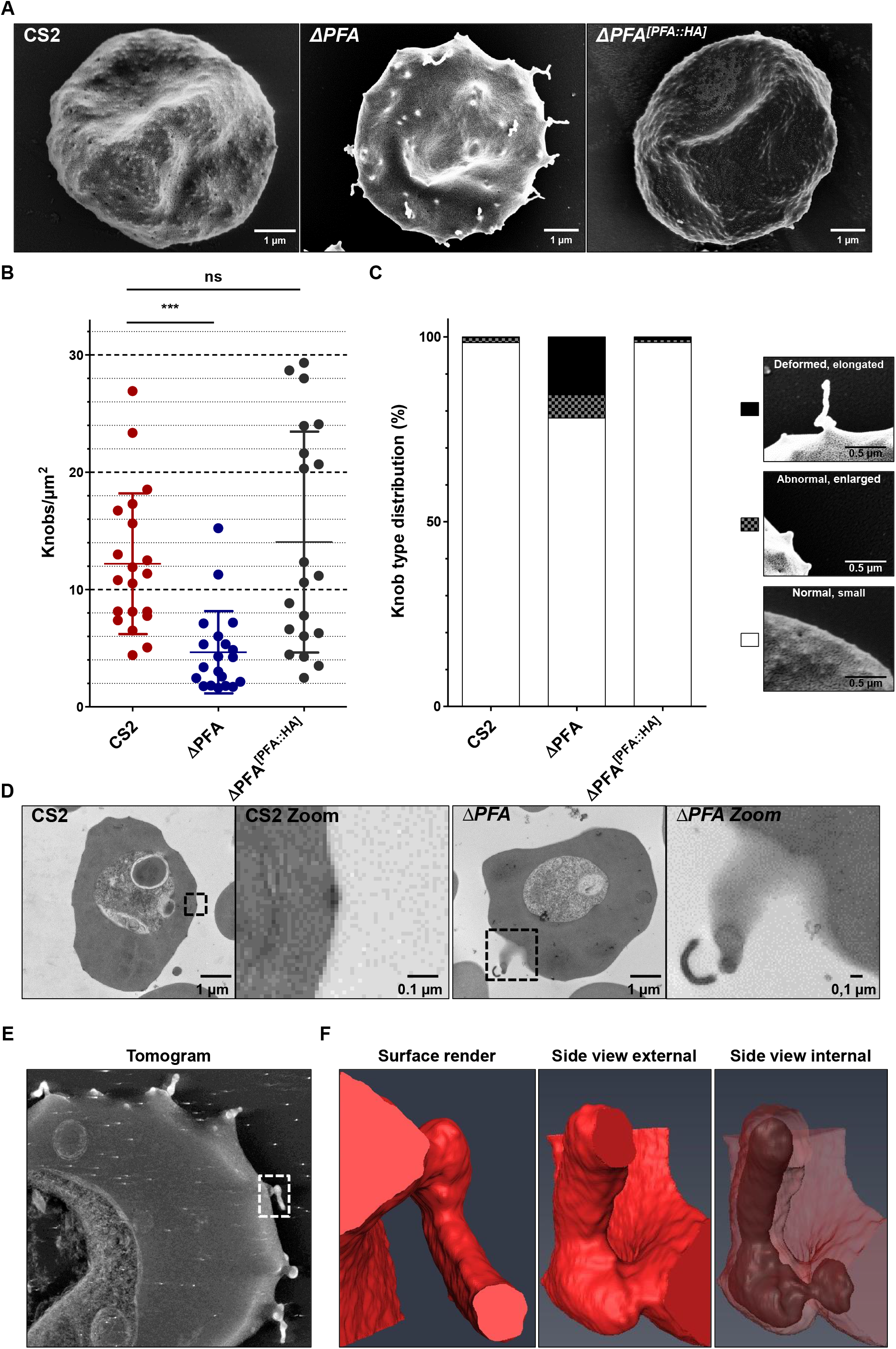
Electron microscopy reveals deformed knob morphologies of Δ*PFA* iRBCs. A) Scanning electron microscopy shows knobs on the surface of iRBCs. The mutant phenotype of Δ*PFA* is alleviated upon reintroduction of episomally expressed PFA66 in Δ*PFA^[PFA::HA]^.* More pictures can be found in Figure S2. B). Quantification of knob density via ImageJ in SEM pictures (n = 20) shows significantly fewer knobs on Δ*PFA* iRBCs. Knob density is restored in the complementation cell line *CS2ΔPFA^[PFA::HA]^*. C) Quantification of knob morphology across all iRBCs. Knobs were grouped into three categories: small knobs, enlarged knobs, and elongated knobs. Then every knob on 20 SEM pictures of the three strains was assigned to one of these categories. Each bar represents the distribution of these knobs in the three categories across all pictures of a strain. The Δ*PFA* strains display an increase in deformed and enlarged knob morphologies compared to CS2 and Δ*PFA^[PFA::HA]^*. D) Internal view of the deformed knobs/mentula via transmission electron microscopy of thin slices. More pictures can be found in Figure S3, E. Electron tomography reveals electron-dense material at the base and interior of deformed knobs/mentulae. The marked area denotes the structure shown in F. F) 3D segmentation of discrete densities within a deformed *knob/mentula* depicted with electron tomography. The example shows a severely deformed knob/mentula. Additionally electron dense material was detected at the base and inside of these structures.

Transmission electron microscopy (TEM) on thin sections prepared from RBCs infected with either WT CS2 or Δ*PFA* parasites substantiated our observations. CS2 parasites produced clearly defined electron-dense knob structures of a restricted diameter and height above the erythrocyte plasma membrane, whereas Δ*PFA* displayed *mentulae* extending from the red blood cell surface into the external medium (Figure 2D, S3). The lumen of these *mentulae* was often extremely electron dense, hinting at their molecular relation to knobs. Occasionally we observed membrane-bound structures extending from RBCs infected with CS2 parasites, but the lumen of these structures was not electron dense; therefore, these structures cannot be classed as *mentulae*. Although not highly abundant in either sample, the morphology of Maurer’s clefts appeared comparable in both samples (data not shown).

### Electron-dense material is a scaffold for mentula structure

Intrigued by the apparent electron-dense core of *mentulae* in TEM, we conducted electron tomography analysis on thick sections prepared from Δ*PFA*-infected erythrocytes. Subsequent 3D reconstruction and surface rendering of the distinct densities in tomographic volumes allows a high-fidelity glimpse into the fine structure of *mentulae*. This analysis verified the presence of electron-dense material within and at the base of *mentulae* (Figure 2E). This material fills out the entire *mentula* rather than only lining the structure, and the distribution of this material closely matched the structure of the *mentula*. In some cases (3 out of 10), the electron-dense material at the base of the *mentula* was connected via a thin bridge to the material inside of the *mentula* (Figure 2F, Supplementary Video 1).

### KAHRP distribution is changed in ΔPFA-infected erythrocytes

KAHRP has long been held to be a crucial knob-associated protein as parasites lacking this protein no longer form knobs^25,26^, and KAHRP truncations show varying aberrant knob phenotypes^27^. For this reason we investigated the localisation of KAHRP in RBCs infected with our Δ*PFA* cell line. Indirect immunofluorescence (IFA) on fixed cells demonstrated a punctate distribution of KAHRP in cells infected with both wild type and mutant cell lines (Figure 3A). Automated analysis with a self-generated ImageJ plugin revealed that KAHRP +ve structures were equally numerous but statistically larger in diameter in RBCs infected with Δ*PFA* than with CS2 (Figure 3B, C). Localisation of other exported parasite proteins via IFA showed no significant difference between the cell lines (Figure S4). To exclude that the KAHRP result was due to non-specific binding of antibodies, for example, we episomally expressed a KAHRP::mCherry fusion in both wild-type and Δ*PFA* parasites. Surprisingly, considering the relatively low resolution of live cell epifluorescence microscopy, (but preserving membrane integrity and cell morphology), KAHRP +ve structures could be seen to emerge from the surface of RBCs infected with Δ*PFA*, possibly representing *mentulae* (Figure 3D, Supplementary Videos 2A-D). Based on the previous result, we wanted to understand whether KAHRP is directly associated with *mentulae* and used immunogold labelling of thin sections derived from CS2 and Δ*PFA* iRBCs to localise KAHRP. Although we encountered high background staining of the RBC cytosol in both cases, analysis revealed considerably more label was associated with knobs/*mentulae* than with the cytosolic background (Figure 3E, F). Analysis of the location of the KAHRP label in relation to the length of the *knob/mentula* or the closeness to the RBC membrane (RBCM) showed no difference between WT and mutant, or between knob/*mentula* morphologies (Figure S5A, B). To gain more insight into the nature of the KAHRP +ve structures, we paired membrane shearing with immunolabelling and STED (stimulated emission depletion^28^) microscopy to study the nature of the KAHRP +ve structures observed above. RBCs infected with CS2 or Δ*PFA* were allowed to bind to a cover slip, hypotonically lysed to obtain access to the internal leaflet of the RBC plasma membrane, fixed, immunodecorated using an α-KAHRP antibody, and imaged via STED microscopy. This technique allows super-resolution visualisation of KAHRP +ve structures from the luminal side of the erythrocyte plasma membrane and can be used to monitor the assembly of KAHRP into knobs (and in this case, *mentulae*). This analysis revealed a punctuate distribution of KAHRP beneath the membrane of RBCs infected with both wild type and mutant Δ*PFA* cells. Although the data initially suggests that there are more KAHRP +ve structures in membranes from Δ*PFA*, automated counting and measurement shows that this impression is likely to be imparted due to a higher mean size of KAHRP +ve objects rather than an actual increase in the absolute number of objects (Figure 3G, H, I). KAHRP +ve structures from Δ*PFA*, as well as being larger, also appear by eye to have a lower circularity, although we were not able to substantiate this with image analysis. This might be in part due to the differences in morphology between knobs and *mentulae*.

**Figure 3.**
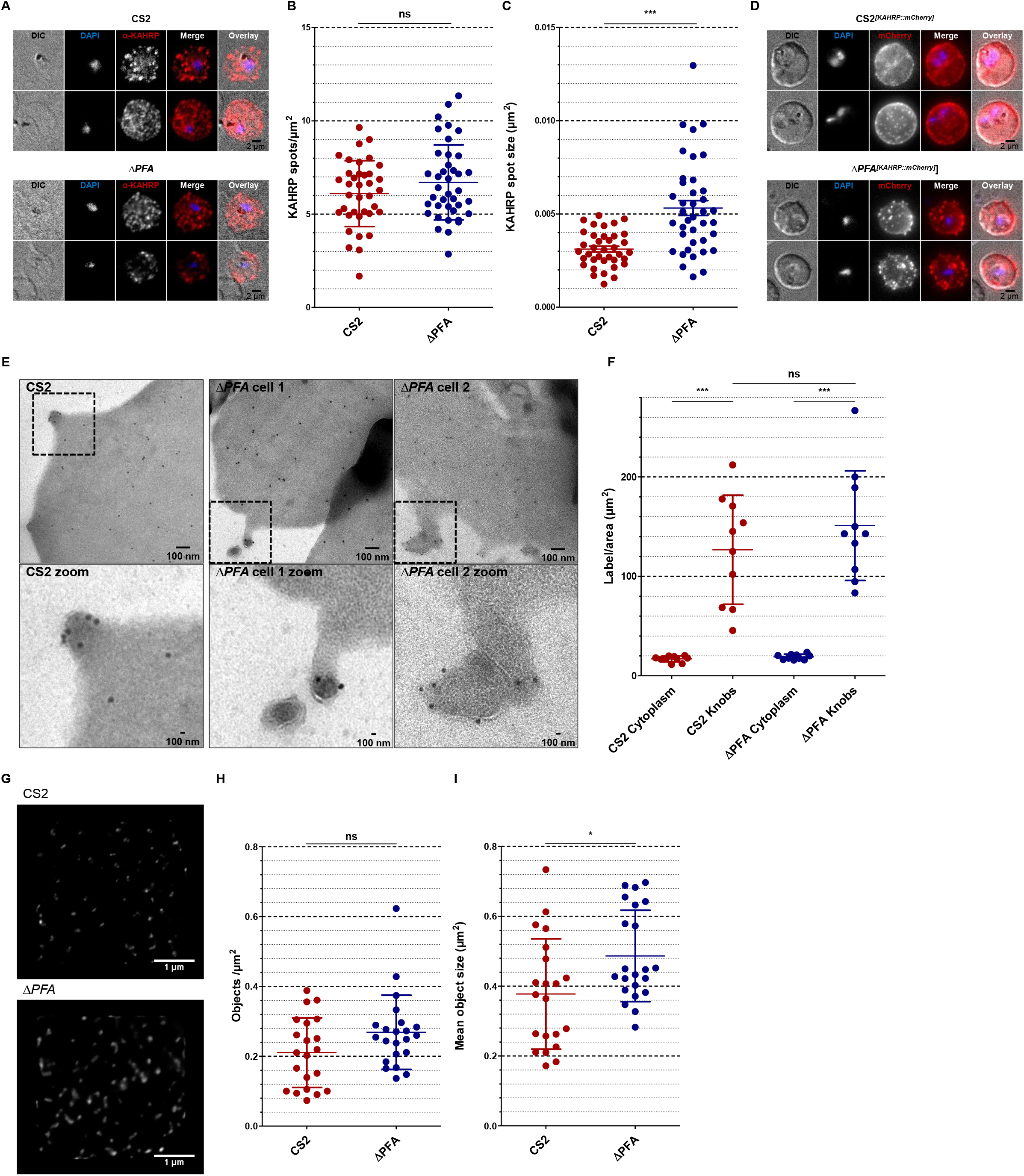
A) Immunofluorescence assay of MeOH-Ac-fixed CS2 and Δ*PFA* using α-KAHRP antibodies reveals punctate patterns. A trend was noticed towards bigger spots in the truncation strain and verified using automated measuring via an ImageJ algorithm (See Figure 3 B, C). D) Live cell imaging of DAPI stained *CS2^[KAHRP::mCherry]^* and Δ*PFA^[KAHRP::mCherry]^.* KAHRP::mCherry can be seen in both cell lines as punctate patterns; however, CS2 displays smaller and more dots. E) Immunogold labelling of iRBC sections in TEM using α-KAHRP antibodies. Images demonstrate label associated with normal knobs and deformed knobs in CS2 and Δ*PFA*, respectively. Framed areas can be seen enlarged below. F) Analysis of label density associated with the cytoplasm and area surrounding knobs. Label density is significantly higher in the area surrounding knobs than the cytoplasm for both strains. G) STED imaging of the KAHRP associated with the internal RBC cytoskeleton. For this analysis CS2 and Δ*PFA* iRBCs were bound to a dish and then lysed hypotonically. The cell body was then washed away, and the remaining cytoskeleton remained as it would be seen from the inside of the iRBC. These samples were then interrogated with an α-KAHRP antibody and STED imaging. G) Representative images of the KAHRP patterns observed in STED from the CS2 and Δ*PFA* cell line. KAHRP signals were often found to be bigger in the truncation cell line. H) Computational analysis of KAHRP signals through a self-made ImageJ tool revealed no difference in KAHRP spot numbers between both cell lines. I) Investigation of mean object size demonstrated a slight increase of KAHRP spot size in Δ*PFA*.

### Mentulae *contain ring/tubelike KAHRP structures but only small amounts of actin*

To further investigate the relation of KAHRP with *mentula* we used rSTED (rescue-stimulated emission depletion^29^) to image both RBCs infected with CS2 or Δ*PFA* parasites episomally expressing KAHRP::mCherry as above (Figure S5C). Infected cells were treated with RFP booster to amplify the mCherry signal, fluorescently labelled wheat germ agglutinin (WGA) to label the RBC glycocalyx (delineating the RBC membrane, RBCM), and phalloidin to stain host cell actin (Figure 4).

**Figure 4.**
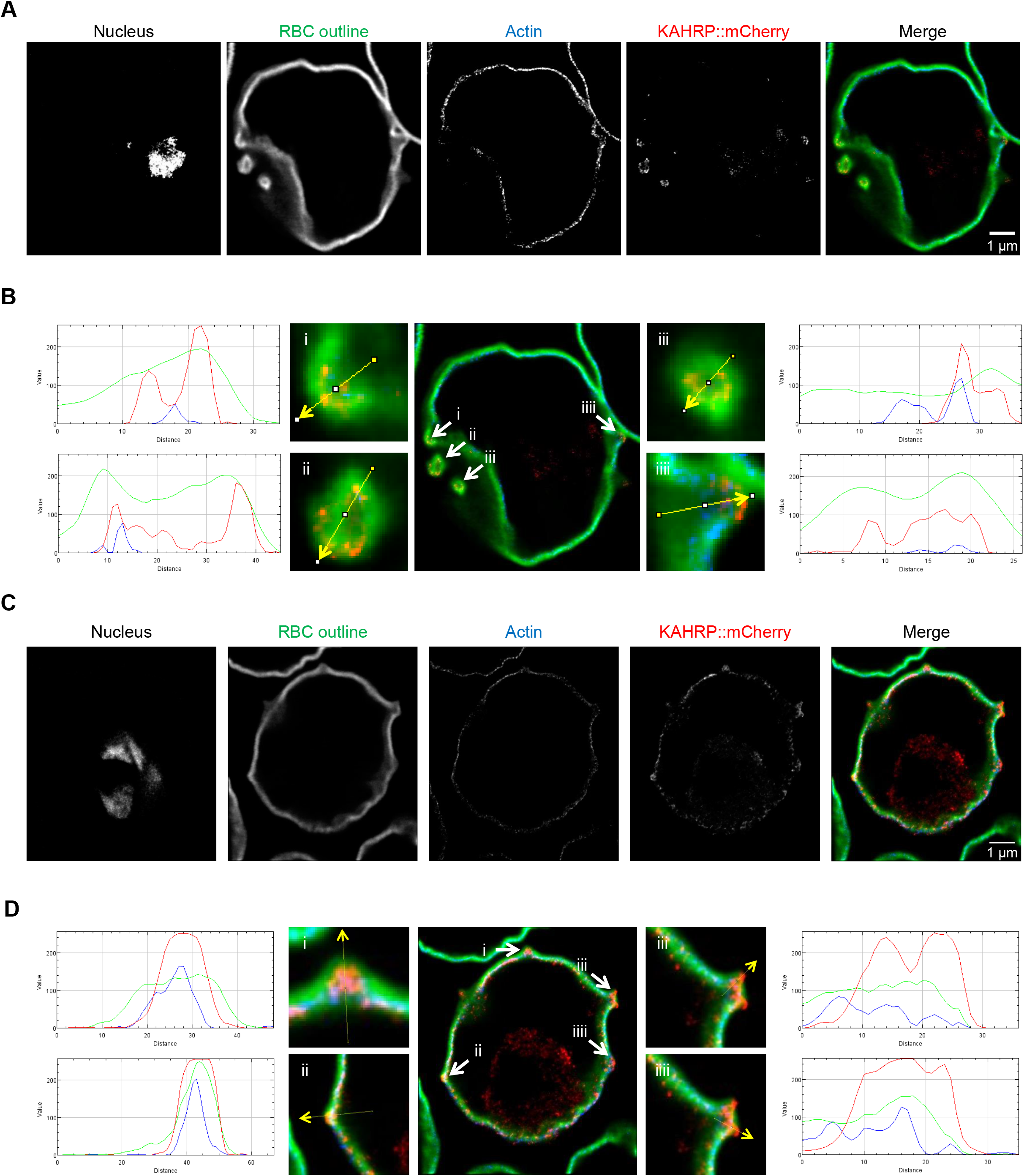
Investigation of the subcellular composition of deformed knobs in Δ*PFA^[KAHRP::mCherry]^* via rSTED imaging in an immunofluorescence assay. A, C) DNA was stained using DAPI; WGA was used to stain the RBC glycocalyx; phalloidin was used to stain actin; and RFP booster was used to label KAHRP::mCherry. B, D) RGB profiles of selected, KAHRP_mCherry-rich structures (likely representing knobs). The RGB profiles demonstrate that in the vertical view, phalloidin (*i.e*. actin) is localized toward the cytosol from the KAHRP structures. The horizontal view shows that the KAHRP-containing structures form a ring structure. These might contain low amounts of actin but are likely filled with other material(s).

Line scan analysis reveals that *mentulae* are bounded by the RBCM. In most cases, KAHRP is found between the RBCM and the actin cytoskeleton, and the KAHRP signal also extends into the central cavity of the *mentulae.* Distribution of host actin closely follows that of WGA, apart from at the base of *mentulae* where it follows a path below the KAHRP staining. Fortuitous sectioning of several membrane-bounded *mentulae* extending from the RBC reveals a ring of KAHRP staining lining the luminal face of the *mentula.* Phalloidin staining was absent in this case, although it was associated with a shorter *mentula* (Figure 4, see Figure S5D for CS2 cell line).

### Chelation of membrane cholesterol but not actin depolymerisation or glycocalyx degradation causes reversion of the mutant phenotype in ΔPFA

A number of lipid- or protein-dependent mechanisms can induce the initiation of curvature of biological membranes and stabilisation of the resulting structures^30^. The *mentulae* we observe here extend up to 0.7 μm from the surface of the red blood cell and appear stable enough that we were able to observe them in live cell imaging. To understand how such structures can be generated, we treated erythrocytes infected with Δ*PFA* parasites with cytochalasin-D (cyto-D) to depolymerise actin, methyl-β-cyclodextrin (MBCD) to chelate membrane cholesterol, or the glucosidases hyaluronidase (HA) and neuraminidase (NA) to degrade RBC glycocalyx. Following treatment, cells were fixed and prepared for SEM, followed by image acquisition and analysis. Neither cyto-D nor HA/NA treatment caused a statistically significant reduction in the number of *mentulae* in cells infected with Δ*PFA* (Figure S6A, B, C, D, E). Treatment with MBCD, while not causing a total reversion of *mentulae* to knobs, did cause a statistically significant alteration in the observed type of *mentulae*, with a decrease in the number of deformed, elongated *mentulae*, and an increase in abnormal, enlarged *mentulae*.

### PFA0660w truncation results in negligible cytoadhesion and aberrant PfEMP1 presentation

As previously mentioned, mature stage infected *P. falciparum* iRBCs develop additional adhesive capabilities to cells of the host, and this process underlies *P. falciparum* pathogenicity^1^. IRBC cytoadherence to purified ligands can be assessed with a binding assay^18^. Prior to genetic manipulation we selected our parasite population for expression of the *var2CSA* variant of *Pf*EMP1 by repeatedly binding them to chondroitin sulphate-A (CSA)^19^. We chose to use the CS2 strain of *P. falciparum* as this strain stably expresses *var2CSA*^18^. A static cytoadherence assay against immobilised CSA demonstrated that, in direct comparison to the parental CS2 strain, Δ*PFA* exhibits massively reduced levels of binding (Figure 5A). Although all experiments were carried out on parasites that had not been maintained in culture for extended time periods (to avoid switching to another var/*Pf*EMP1 variant), we wanted to verify that Δ*PFA* still expressed *Pf*EMP1^var2csa^. We therefore used flow cytometry on intact iRBCs using antisera specific against VAR2CSA (a kind gift of Benoît Gamain) to verify VAR2CSA expression and surface exposure. On erythrocytes infected with Δ*PFA*, surface expression of VAR2CSA was reduced 60% compared to wild type parasites (Figure 5B, Figure S6F for histograms). Immunofluorescence (IFA) using the same antisera on fixed cells demonstrated that both CS2 and Δ*PFA* express VAR2CSA to similar levels, with punctuate staining distributed across the host cell. A control IFA on cells infected with 3D7 strain parasites that had not been selected for VAR2CSA expression showed only background fluorescence, verifying the specificity of the result (Figure S6G).

**Figure 5.**
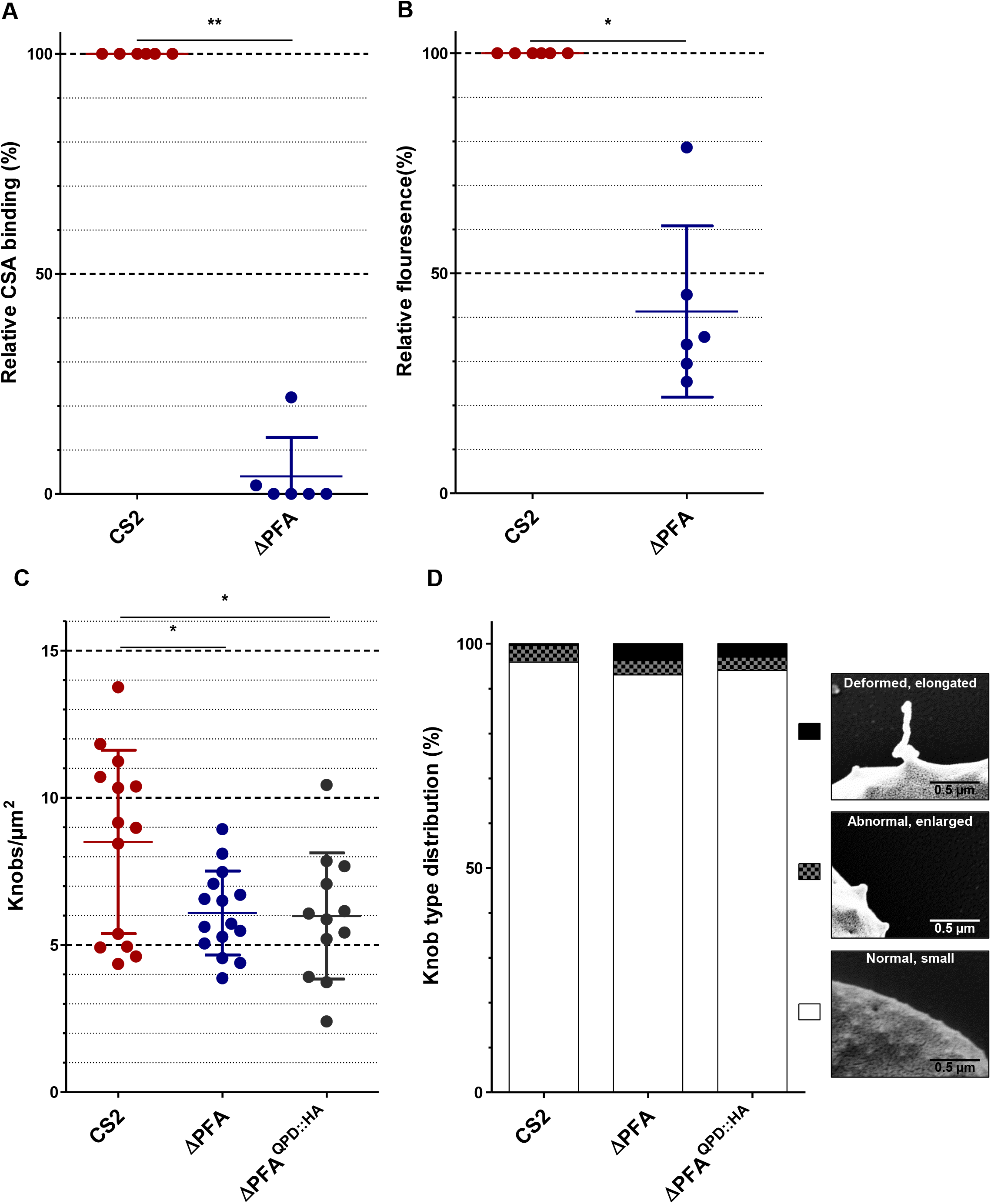
A) Δ*PFA* displays negligible cytoadherence and lower *Pf*EMP1 surface exposure than CS2. CS2 and Δ*PFA* were assayed to test their ability to adhere to immobilized CSA in Petri dishes using microscopic counting of the cells. Cytoadhesion strength is expressed relative to CS2. Results are shown for six binding assays. B) Analysis of *Pf*EMP1 surface exposure via flow cytometry. IRBCs were stained with DAPI and αVAR2CSA antiserum followed by a Cy3-coupled secondary antibody. Δ*PFAs* have lower *Pf*EMP1 surface exposure than CS2 in six independent experiments. C) Expression of a PFA variant featuring a mutated HPD motif in the cell line Δ*PFA^[QPD::HA]^* does not complement reduction in knob abundance (C) and knob deformation (D) observed in Δ*PFA*. As in previous experiments, iRBCs off the three cell lines were purified and imaged via SEM. Knobs were then counted and grouped into three categories.

### Complementation of PFA66 function requires an intact J-domain

HSP40s such as PFA66 are known interactors and regulators of HSP70s^11,12,31^, and HSP70-independent functions of HSP40s are rare. The J-domain of HSP40s is crucial both for recruitment of the partner HSP70 and stimulating the ATPase activity of the HSP70 partner^31^. Having shown above that we can achieve functional complementation via episomal expression of a full-length copy of PFA66, we were interested to know whether a functional J-domain (and hence a functional interaction with a HSP70) is required for phenotypic complementation. To this end, we expressed in the Δ*PFA* parasite line a full-length copy of PFA66 with a H111Q amino acid replacement converting the HPD motif of the J-domain into the non-functional QPD sequence (Figure S6H) and assayed *knob/mentula* morphology and density as a surrogate for all other phenotypic assays. While expression of full-length PFA66 was able to revert the mutant knob phenotype (Figure 2B, C), similar expression of the non-functional full-length protein failed to complement the wild-type phenotype (Figure 5C, D) in either density or morphology of knobs/*mentulae*.

## Discussion

Although it has long been recognised that malaria parasites export a substantial number of proteins to their host cell, the mature human erythrocyte, the function of many of these proteins remains unknown^6,7,10^. *P. falciparum* exports a larger number of proteins to the host cell than related species, and one family that is highly represented amongst this expansion is that of J-domain proteins (Hsp40s)^8^. A previous medium-throughput study identified the exported Type II Hsp40 PFA66 as likely to be essential and resistant to inactivation via double-crossover integration^9^. In this study, we have used selection-linked integration (SLI)^17^ to generate parasites expressing a severely truncated form of PFA66. This strategy deletes the entire substrate-binding domain of the expressed protein, resulting in a non-functional truncation mutant that is incapable of carrying out its biological function, and may well exhibit a dominant negative effect. The resulting parasites are highly viable but exhibit severe abnormalities in host cell modification.

Parasite growth following truncation of PFA66 was slightly higher than that of wild type CS2 parasites, this effect only becoming evident after four growth cycles. A similar effect was previously observed in a Δ*GEXP07* parasite line; however, the significance of this result remains unclear as the reduction in metabolic burden by the loss of only one gene/protein is likely to be negligible^32^. Lysis of infected erythrocytes in a sorbitol uptake assay was also slightly reduced, implying either a reduction in novel permeation pathway activity, or potentially an increase in host cell stability via mechanical means. As a reduced NPP activity would be expected to lead to slower, not faster, parasite growth, we interpret this to be the result of increased robustness of the erythrocyte plasma membrane through an unknown mechanism.

The most striking result of our study was the observation that, in the Δ*PFA66* cell line, normal knob biogenesis was significantly inhibited with regard to both knob density and morphology. Although earlier knockout studies have observed a reduction in knob formation, or slight alteration in knob morphology, to our knowledge ours is the first study to demonstrate such a dramatic alteration in knob structure upon inactivation of a single gene^9,32^. Indeed, so different are the structures we observe to classic knobs that we suggest calling them *mentula* to distinguish them from the ‘normal’ surface extensions. *Mentulae* differ from knobs both in their size/length, reaching up to 0.7μm from the erythrocyte surface. Additionally, *mentulae* that split into separate branches can be observed. KAHRP, a protein known to be required for correct knob formation, can be localised to *mentulae*, although its distribution (based on live cell imaging, immunofluorescence, and membrane shearing paired with STED microscopy) seems to be different from that observed in cells infected with wild type parasites. Several studies suggest that KAHRP is integrated into higher order assemblies during parasite development, eventually resulting in a ring structure underlying the knob^33,34^. Deletions in specific KAHRP domains lead to less incorporation into such structures and also appear to influence the generation of sub-knob spiral structures of unknown molecular composition^34^. The possibility exists that KAHRP, while being necessary for correct knob formation, is itself not the major structure-giving component but merely serves as a scaffold for assembly of further higher order molecular structures, which themselves generate the necessary vector force to allow membrane curvature and push the knobs above the surface of the erythrocyte. KAHRP has been proposed to be especially vulnerable to misfolding due to its unusual amino acid composition, which would make it a likely client for chaperones / cochaperone systems^14^. Following this logic, if PFA66 is involved in the assembly of KAHRP into higher order assemblies, interruption of this process could cause knock-on misassembly of the spiral and therefore aberrant knob structures. Although we were not able to visualise spiral structures at the base of *mentulae*, our observed phenotype could be explained by runaway lengthening of such a spiral structure (Figure 6). Indeed, the electron density we observed on tomograms is consistent with a high molecular weight structure as a form-giving scaffold for the formation of *mentulae*. Within the resolution limit of our study, KAHRP itself appears to line the inner leaflet of the membrane-bounded *mentulae* and is thus unlikely to itself be a major component of the electron dense core of the *mentulae.* The action of host actin is known to be required for the generation of knobs^34^; however, although we could successfully visualise actin below the erythrocyte plasma membrane, it appears to be largely excluded from *mentulae* and thus cannot be responsible for maintaining the form of these structures, a view supported by the lack of action of cyto-D on *mentula* structure. Similarly, enzymatic removal of the erythrocyte glycocalyx had no effect on *mentulae*, excluding a role for this in membrane shaping. Chelation of membrane cholesterol via treatment with MBCD did cause change in the morphology of *mentulae*, but not a complete reversion to normal knob structures. Removal of cholesterol from biological membranes has been observed to cause an increase in membrane stiffness^35,36^, and this may explain the morphological reversion upon MBCD treatment, with a stiffer membrane being more resistant to the pushing force within the *mentulae*. Alternatively, removal of cholesterol and subsequent breakdown of so-called lipid rafts may interfere with the higher order organisation of membrane-bound factors involved either directly in membrane curvature or required for coordination of proteins internal to the *mentulae* that generate force. Taken together, our data strongly implies that both knobs and *mentulae* contain a so far unknown component, likely proteinaceous, which is required for force generation, subsequent membrane curvature, and resulting morphology. Interruption of correct assembly of this factor leads to aberrations in knob formation, eventually leading to the appearance of *mentulae*. Our, and others’, data implicates KAHRP as having a role in this process, but it is unlikely to be the only structural protein involved.

**Figure 6.**
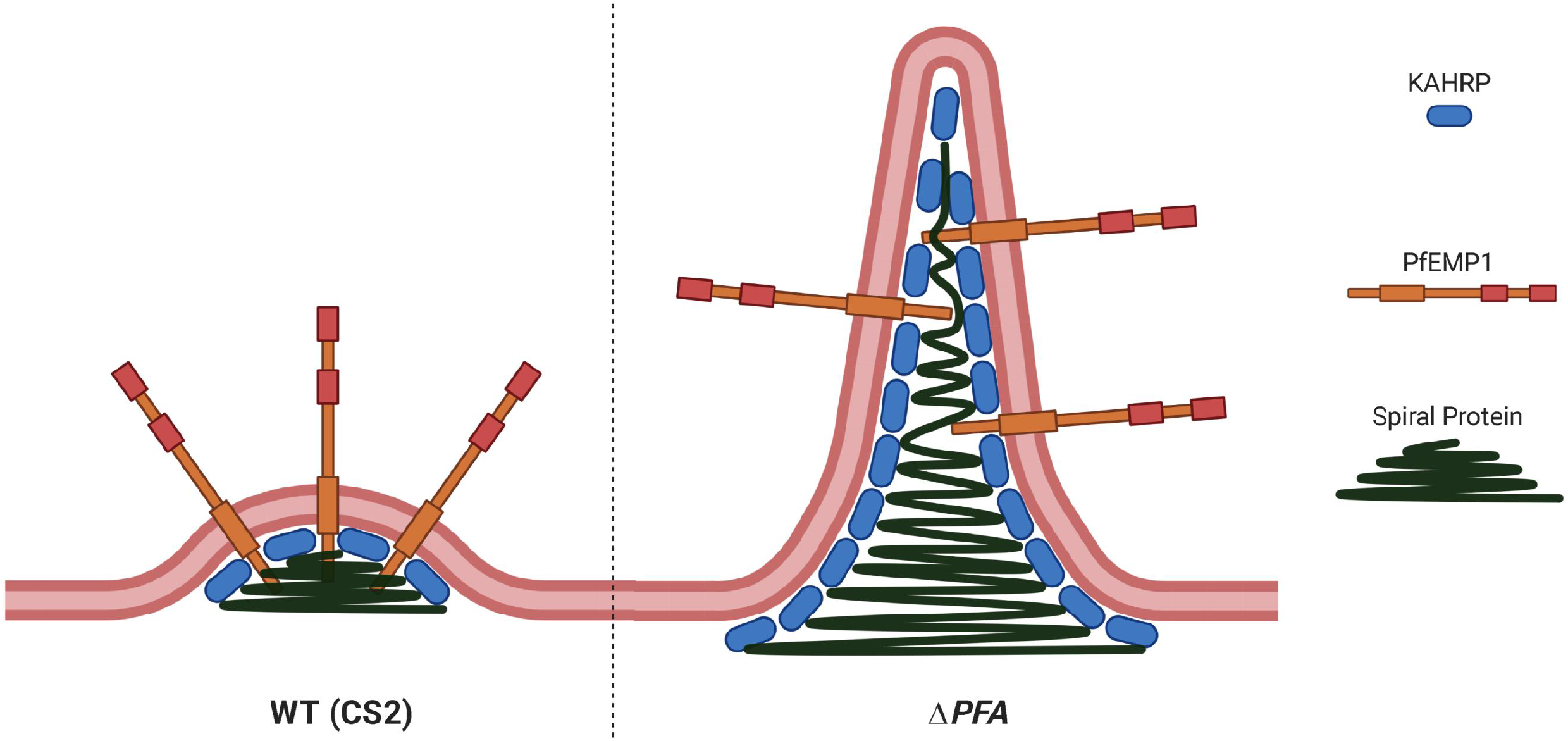
Proposed model for *mentula* bioformation. In opposition to normal knob formation in the CS2 cell line (left) runaway extension of the spiral underlying *mentulae* in Δ*PFA* could drive their elongation (right). KAHRP is still present and associated with the inner lumen of *mentulae*, PfEMP1 anchored in the *mentulae* is incorrectly presented and has thus a reduced cytoadherance capacity.

In other systems, HSP40s, through their role as a regulator of HSP70 chaperone activity, have been shown to have a role in both assembly and disassembly of protein complexes^37^, and it is tempting to suggest that the phenotype we observe here is due to incorrect complex assembly. Alternatively, PFA66 may be required for the correct transport of accessory proteins required for complex formation such as those proposed in a recent study^34^, and thus play an indirect role in correct assembly of high-molecular weight complexes. In support of this hypothesis, PFA66 is known to associate with J-dots, highly mobile structures within the infected erythrocyte that are also known to contain a number of HSP70s^13,15,16,20^. It is also feasible that PFA66 is required for the disassembly of incorrectly folded or assembled knob protein complexes and that our knockout reveals so far unknown quality control mechanisms.

Knobs are required for correct presentation of the major virulence factor *Pf*EMP1, and high affinity binding of such to endothelial receptors^38^. Although previous studies suggested that KAHRP and *Pf*EMP1 formed a ‘precytoadherance complex’ at the Maurer’s clefts^39^, later evidence suggest strongly that *Pf*EMP1 is incorporated only into knobs once they have been at least partly formed^34,40^. Erythrocytes infected with Δ*PFA66* parasites showed a 63% reduction in *knob/mentulae* density and a 10-fold increase in the frequency of abnormal knob phenotype, and we observed an almost total lack of cytoadherence in iRBC infected with Δ*PFA66* parasites. This data strongly supports the view that, even in the normal knobs present, less *Pf*EMP1 was correctly presented and could take part in cytoadherence. Flow cytometry determined an almost 60% drop in cell surface recognition of the VAR2CSA variant of *Pf*EMP1, although immunofluorescence suggests that both cell lines express similar amounts of this protein. The total loss of cytoadherence may be the consequence of several distinct factors: a) less total surface *Pf*EMP1, b) fewer knobs with correctly loaded *Pf*EMP1, and c) a significant number of aberrant knobs/*mentulae*. Moreover, we cannot exclude that *Pf*EMP1, which needs to be correctly folded to bind specific receptors, does not assume the correct tertiary structure due to the lack of the necessary chaperone/co-chaperone system.

As previously mentioned, HSP40s generally act in concert with members of the HSP70 family^12^. The erythrocyte is known to contain significant amounts of residual human HSP70s^41^ and additionally a parasite-encoded HSP70, *Pf*HSP70-X ^15^. We have previously demonstrated that a knockout of *Pf*HSP70-X leads to a reduction in virulence characteristics, including cytoadherence^42^. Significantly, however, iRBCs infected with Δ*70-x* parasites were covered with normal knob structures at a density comparable to that of the wild type^42^. Hence, although both knockouts showed defects in virulence functions, the mutant phenotype of iRBCs infected with Δ*PFA66* is distinct and significantly more dramatic than that in iRBCs infected with Δ*70-x* parasites. PFA66 has been reported to undergo a functional interaction with *Pf*HSP70-X, HsHSP70, and HsHSC70^43,44^. Considering the difference in phenotype between Δ*70-x* and Δ*PFA66*, here we must conclude that, if a HSP70 is involved, it is more likely to be of human rather than parasite origin. We cannot formally exclude that PFA66 functionally interacts with both human and parasite HSP70 homologues and that the phenotype we observe is a combination of the effect on both chaperones, but the balance of probabilities suggests that morphological abnormalities observed in this current study are largely due to an interruption of functional PFA66-*Hs*HSP70/HSC70 interactions.

The J-domain of HSP40s contains a characteristic HPD motif that is required both for binding and ATPase activation of partner HSP70s^37^. As the unlikely possibility existed that our observed phenotype was due to a HSP70-independent function of PFA66, we carried out complementation analysis with a full-length copy of PFA66 expressed from an episome, or a copy containing a H111Q mutation that renders the J-domain inactive. As only the wild type protein was able to complement the mutant knob phenotype, we conclude that the effects we observed upon PFA66 truncation are due to either a dominant-negative effect of the truncated protein on proper functioning of the *Hs*HSP70 chaperone system within the host erythrocyte, or an effect due to lack of co-chaperone activity via deletion of an essential functional domain (SBD) of the HSP40. Either way the results support *Hs*HSP70 involvement. A potential role for residual human HSP70 in host cell modification and parasite virulence has been suggested for almost 20 years^45^, but to our knowledge our current study is the first to provide strong experimental evidence implicating human HSP70s in these processes.

To conclude, in this study we show data suggesting that correct biogenesis of knobs in malaria-infected erythrocytes is a complex process necessitating a number of proteins, the molecular identity of some of which remains enigmatic. Our data suggests that KAHRP, while obviously required for knob generation, may not directly provide a scaffold for knob structure. More importantly, our data also reveals that residual human HSP70 within the infected erythrocyte is involved in parasite-driven host cell modification processes. To our knowledge, this is the first time a host cell protein has been implicated in parasite virulence, and this observation opens up exciting new avenues for the development of new anti-malarials.

## Materials & Methods

### Vector construction

The ~1kbp PFA66 targeting region was amplified using the primers PFA_NotI_F and PFA_MluI_R and cloned into pSLI^TGD^ using NotI-HF and MluI (NEB)^17^ (kind gift of Tobias Spielmann). The complementation plasmid PFA::HA was generated by excising the *PFA0660w* coding sequence and promoter from the plasmid pAD-A660-GFP^13^ with the restriction enzymes NotI-HF and BssHII and cloning them into pARL-PFF1415c-3xHA (BSD, a kind gift of Sarah Charnaud). The resulting plasmid was subsequently used to generate the QPD::HA plasmid via QuickChange PCR using PFA_Quick_QPD_F and PFA_Quick_QPD_R primers. Upon verification of the QPD mutation, the insert consisting of the *PFA0660w* promoter and coding sequence was re-cloned into the same vector to avoid mutation due to the PCR step. The KAHRP::mCherry plasmid was generated by amplifying mCherry with the primers mCherry_AvrII_F and mCherry_XmaI_R and cloning them into a pre-existing plasmid pARL2_KAHRP containing the native KAHRP promoter and KAHRP coding sequence (kind gift of Cecilia Sanchez). All primers are listed in Supplementary Table 2A.

### Cell culture methods

*P. falciparum* parasites were cultured at 37°C with 90% N2, 5% CO_2_, and 5% O_2_ according to established methods^46^. Parasites were maintained at a haematocrit of ~5% in A^+^ or O^+^ human blood obtained from the blood banks in Marburg and Heidelberg, respectively, and maintained with RPMI1640 (Gibco) containing 200 μM hypoxanthine, 160 μM neomycin (Sigma Aldrich) and 10% human plasma. Parasitaemias were evaluated from smears prepared from the blood cultures, which were fixed in 100% MeOH and stained with ddH_2_O/10% Giemsa solution (Merck). Parasites were transfected with 150 μg of plasmid and treated with 2.5nM WR99210 (HS lines) or 12μg/ml blasticidin (Invivo Gen). Transfectants were propagated in fresh O^-^ blood using RPMI1640 (Gibco) with 5% human plasma, 5% albumax II (Invitrogen) and other additives as above until parasites re-appeared^47^. Selection-linked integration was performed according to Birnbaum et al.^17^. Briefly, following reappearance after the initial transfection, parasites were treated with 400 μg/ml G418 (Thermo Fisher Scientific) until resistant parasites were observed. Parasites were synchronized before experiments using sorbitol-induced lysis^48^. For this mixed-culture parasites were incubated in 5% sorbitol for 10 min, washed with parasite culture medium and re-cultivated. Routine selection for CSA-binding parasites was performed according to standard protocols^19^. Late-stage parasites were enriched via gelatine flotation for 1 hr / 37 °C ^49^. Subsequently parasites were resuspended in cytoadhesion media (pH 7.2, prepared from RPMI1640 powder (Life technologies)) and incubated in cell culture flasks pre-treated with CSA (PBS pH 7.2/1 mg/ml CSA overnight / 16 °C and blocked with PBS pH 7.2 / 1% BSA) for 1 hr / 37 °C. After careful washing with cytoadhesion media, the remaining bound parasites were washed off and re-seeded.

### Chemical or enzymatic treatment of iRBCs

For these experiments, magnetically purified iRBCs (~1 x 10^7^ per condition) were used. Cytochalasin-D treatment was carried out with RPMI1640 / 10μm cyto-D for 10 min at RT, incubation with RPMI1640/10 mM MBCD was performed for 20 min at 37 °C, RPMI1640/ 30mU neuraminidase or RPMI1640/30U hyaluronidase treatments were performed for 1 hr at 37 °C. Following treatment, samples were processed for SEM.

### MACS purification

For some protocols, late-stage parasites were magnetically purified using a VARIOMACS with a CS-column^50^. Briefly ~1 ml packed erythrocytes (~10% parasitaemia) were applied to a CS column, washed with PBS / 3% BSA and finally eluted into PBS.

### Microscopy methods

Live cell imaging was performed on DAPI-stained (1 ng/ml) parasites using a Zeiss Axio-Observer microscope and AxioVision software. For immunofluorescence assays, parasites were fixed on microscopy slides using 90% acetone / 10% MeOH for 5 min / −20 °C. Cells were then blocked using PBS / 3% BSA for 1 hr / RT and incubated in a humid chamber overnight with the primary antibody diluted in blocking buffer (for antibodies see Supplementary Table 2B). On the next day, PBS-washed slides were treated with the secondary antibody, diluted 1:2,000 in blocking buffer for 2 hr / RT, subsequently washed again, DAPI stained (0.1 ng/ml in PBS), and imaged using a Zeiss Axio-Observer microscope and AxioVision software. RSTED imaging was carried out as recently reported in great detail^51^. Wheat germ agglutinin Alexa Fluor® 488 conjugate (Thermo Fisher) was used according to the supplier’s specification. Phallodin-Atto 647N (Thermo Fisher) was diluted 1:500 in PBS and incubated for 30 min / RT to stain the actin cytoskeleton. RFP booster ATTO594 was used 1:200/2hr/RT to enhance RFP fluorescence. IMSpector imaging software (Abberior Instruments GmbH) was used for image capture and deconvolution of STED images, and AxioVision software was used for other acquisitions. Images were processed using ImageJ. Brightness and contrast was adjusted to reduce background and enhance visibility. No gamma adjustments were applied to any images, and all data is presented in accordance with the recommendations of Rossner and Yamada^52^.

### Protein-based methods

Protein extracts were prepared from 1 x 10^8^ MACS-purified iRBCs. These were resuspended in PBS and boiled in Laemmli loading buffer for 10 min at 99 °C. Soluble fractions were separated via centrifugation (4 °C, 35,000 g) and an equivalent of 1 x 10^7^ parasites loaded onto each well of 12% acrylamide gels. Equinatoxin (EQT) treatment and fractionation of MACS-purified iRBCs was carried out as described by Külzer et al.^22^ but using 4 haemolytic units of EQT at RT for 6 min. Western blot / immunodetection was carried out via semi-dry blotting, blocking in 5% milk powder (1 hr / RT), incubation with primary (overnight / 4 °C), washing three times with PBS, incubation with the secondary (2 hr / RT) antibody in blocking buffer, washing three times with PBS, and visualization via x-ray films. Antibody sources and dilutions can be found in Supplementary Table 2B.

### Membrane shearing

For investigation of the internal structure of the RBC cytoskeleton membrane shearing was employed according to established protocols^34,53^. Briefly, a (3-aminopropyl)triethoxysilane-treated ibidi dish was incubated with 150 μl PBS / 1 mM with Bis(sulfosuccimidyl)suberate for 30 min / RT, washed with PBS, and incubated with 150 μl ddH_2_O / 0.1 mg/ml erythroagglutinating phytohaemagglutinin for 2 hr / RT. Dishes were rinsed three times with PBS and quenched using PBS / 0.1 M glycine for 15 min / RT. Approximately 1 x 10^7^ MACS-purified iRBCs were added and incubated for 3-4 hr, washed, and sheared using 5P8-10 buffer (5 mM Na_2_HPO_4_ / NaH_2_PO_4_, 10 mM NaCl, PH 8), while angling the dish at 20°. Samples were then blocked using 150 μl of PBS / 1% BSA, treated with the primary α-KAHRP antibody overnight / 4 °C in blocking buffer, washed three times, incubated with the secondary α-mouse^ATTO549^ for 1 hr / RT in blocking buffer, and finally washed three times before imaging via rSTED.

### Sorbitol lysis

Assessment of NPP activity was carried out according to Baumeister et al.^23^. For each measurement, 40 μl of 2% trophozoite culture was resuspended in 150 μl lysis buffer (290 mM sorbitol, 5 mM HEPES, pH 7.4) and incubated for 30 min / 37 °C. Remaining RBCs were then pelleted at 1,600 g / 2 min, and the absorbance of the resulting supernatant was measured at OD_570nm_. Samples were then compared to a total lysis control, which was generated in parallel using ddH_2_O instead of lysis buffer.

### Flow cytometry

Infected erythrocytes were fixed for 24 hr at 4 °C using PBS/4% paraformaldehyde/0.0075% glutaraldehyde and stained with DAPI (1 ng/ml) prior to analysis with a BD Canto. In the growth experiments, both cell lines were diluted after every growth cycle with the same factor in order to support parasite growth. Both parasite cell lines were seeded with the same parasitaemia and diluted after every cycle to avoid ‘crashing’ the culture. Parasitaemias were measured before and after every dilution by staining of iRBCs with DAPI and flow cytometry. For staining VAR2CSA on the RBC surface, live parasites were incubated with VAR2CSA antiserum (11P, rabbit, a kind gift of Benoit Gamain) and α-rabbit-Cy3 for 30 min each and then processed for flow analysis as detailed above.

### Cytoadherence

IRBC cytoadhesion to immobilized CSA was investigated using MACS-purified late-stage parasites^18^. Parasites were applied in cytoadhesion media (pH 7.2, made from RPMI1640 powder (Life technologies) to pre-treated spots (PBS pH 7.2/1 mg/ml CSA overnight at 16 °C, blocked with PBS pH 7.2/1% BSA for 1 hr at RT), and then washed with PBS on a Petri dish. After incubation for 1 hr at RT, non-bound parasites were washed away using cytoadhesion medium. Parasites were then fixed using PBS/2% glutaraldehyde for 2 hr at RT and stained with PBS/10% Giemsa for 10 min at RT prior to imaging using a Zeiss Axio Observer microscope and counting with Ilastik^54^ and ImageJ software.

### Electron microscopy

For scanning electron microscopy, purified parasites were fixed using PBS /1% glutaraldehyde for at least 30 min at RT. After washing, parasites were bound to coverslips (pre-treated with 0.1% polylysine for 15 min at RT), washed again, and dehydrated in acetone gradients (ddH_2_O, 25% Ac, 50% Ac, 75% Ac, 100% Ac, 10 min each) followed by critical point dehydration and coating with 5 nm Pd-gold. Cells were imaged using a Zeiss Leo 1530 electron microscope (SE2 detector, ~12,000 x magnification)^55^. For transmission electron microscopy, parasites were fixed in 100 mM Ca-cacodylate/4% paraformaldehyde/ 2% glutaraldehyde, embedded in Spurr and cut into ~70 nm sections. Some samples were fixed using 100 mM Ca-cacodylate/4% paraformaldehyde/ 0.1% glutaraldehyde and treated according to the Tokuyasu protocol for immunogold labelling of KAHRP using an α-KAHRP (rabbit) antibody and a secondary goat α-rabbit-gold conjugated antibody^56^. Some of the EM sections were used without post-contrasting, while some were post-contrasted using 3% uranyl acetate and ddH_2_O/0.15 M Na-citrate/0.08 M Pb(NO_3_)_2_ / 0.16 M NaOH for 2 min. Imaging was performed using a Jeol 1400 microscope operating at 80kV. For electron tomography ~350 nm thick sections were used and examined in a TECNAI F30, 300kV FEG, FEI electron microscope (EMBL Heidelberg). The resulting tomograms were processed using IMOD, ETOMO image/ volume processing software package and the Amira, volume visualisation software.

### Statistics

Statistics were calculated in prism or Excel using unpaired, two-tailed t-tests. p > 0.05 = non-significant (ns); *p < 0.05; **p < 0.01; ***p < 0.00. Figures show mean and standard deviation.

### ImageJ Macro

The ImageJ/Fiji^57^ macro computes the local maxima of each object on the smooth probability map (PM) images generated by ilastik pixel classification. The ilastik ^54^ pixel classification workflow is used to reduce the background in the images and enhance the foreground pixels. To segment each object in the probability map images the local maxima is used as a seed for the 3D watershed plugin. The approach allows to separate close objects and creates masks that are used to measure size and intensity on the raw images. The macro and the instructions on how to use it can be found at: https://github.com/cberri/2D_AutomatedObjectsDetection_ImageJ-Fiji

## Acknowledgements

We wish to thank the blood banks of the University Hospitals in Giessen and Marburg for providing blood. Further we would like to thank the EMCF at the University of Heidelberg (in particular Stefan Hillmer), the EMCF of EMBL Heidelberg (Yannick Schwab and Martin Schorb), the FACS core facility at the ZMBH (Monika Langlotz), as well as the Infectious Diseases Imaging Platform (Vibor Laketa) and Marie Freudenberg. We thank Tim Bostick for proofreading. This work was supported by DFG grant PR1099/8-1 to JMP.

## Abbreviations

SLI: selection-linked integration
(i)RBC: (infected) red blood cell
PV(M): parasitophorous vacuole (membrane)
RBCM: red blood cell membrane
PPM: parasite plasma membrane
BSD: blasticidin
NEO: neomycin
KAHRP: knob-associated histidine-rich protein
SEM: scanning electron microscopy
TEM: transmission electron microscopy
(r)STED: (rescue) stimulated emission depletion
NPP: new permeability pathways
IFA: immunofluorescence assay
EQT: equinatoxin
CSA: chondroitin-sulphate-A
MC: Maurer’s clefts

---

**Figure S1.**
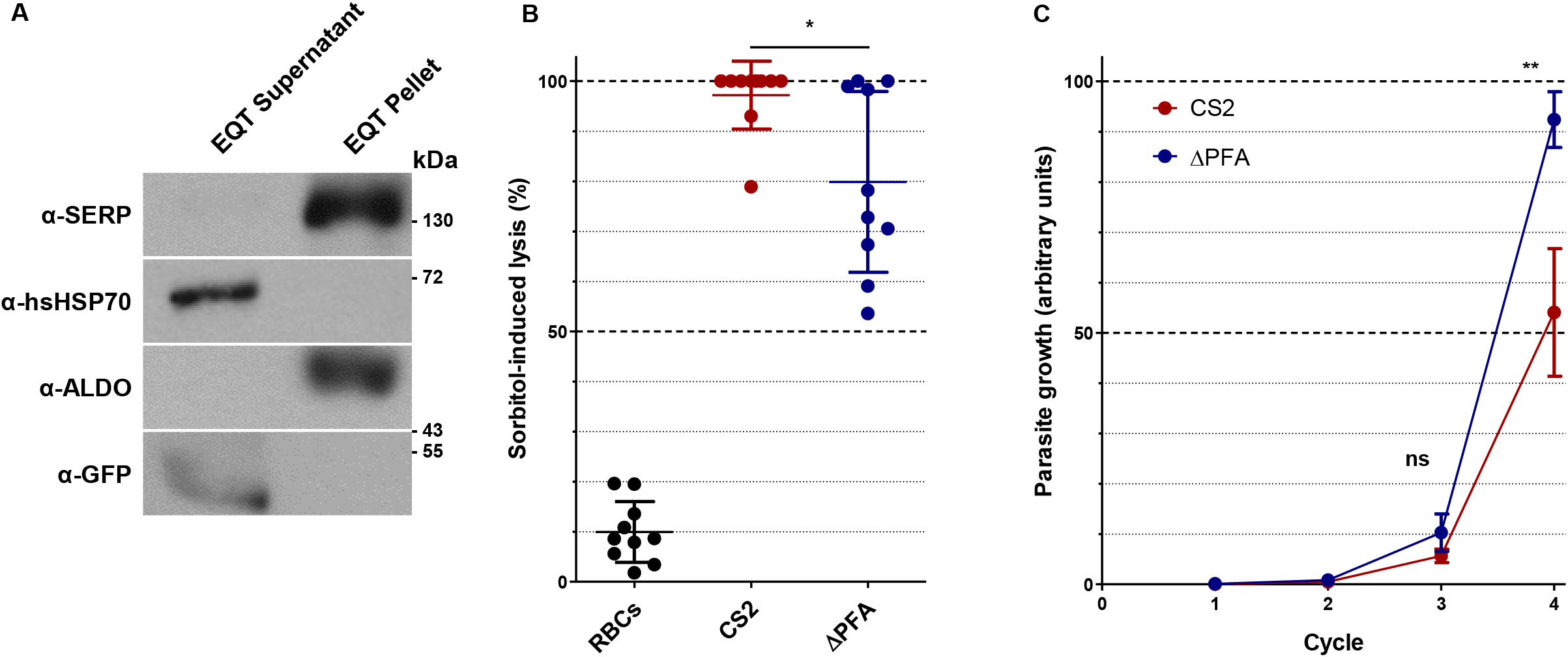
A) An equinatoxin lysis experiment demonstrates export of truncated PFA::GFP. Equinatoxin (EQT) treatment selectively lyses the RBC membrane but leaves the PVM and PPM intact. Consequentially, parasite proteins exported to the host cell are found in the supernatant, while other parasite proteins remain in the pellet. Detection of the PV protein SERP and the parasite protein ALDO in the pellet fraction demonstrates intactness of the PV membrane and PPM, respectively. Truncated PFA::GFP was detected alongside human HSP70 and the exported parasite protein GBP in the supernatant fraction, demonstrating its export to the iRBC. B) Δ*PFA* display a slight decrease in NPP activity when compared to CS2. iRBCs were incubated with the hypotonic agent sorbitol, and NPP activity was assessed by measuring the OD of the supernatant. Results are shown for ten replicates. C) Growth of CS2 and Δ*PFA* was measured over four cycles via flow cytometry of DAPI-stained, fixed parasites. Δ*PFA* show a slight growth advantage over CS2 in the last cycle. Results are shown for three independent experiments.

**Figure S2.**
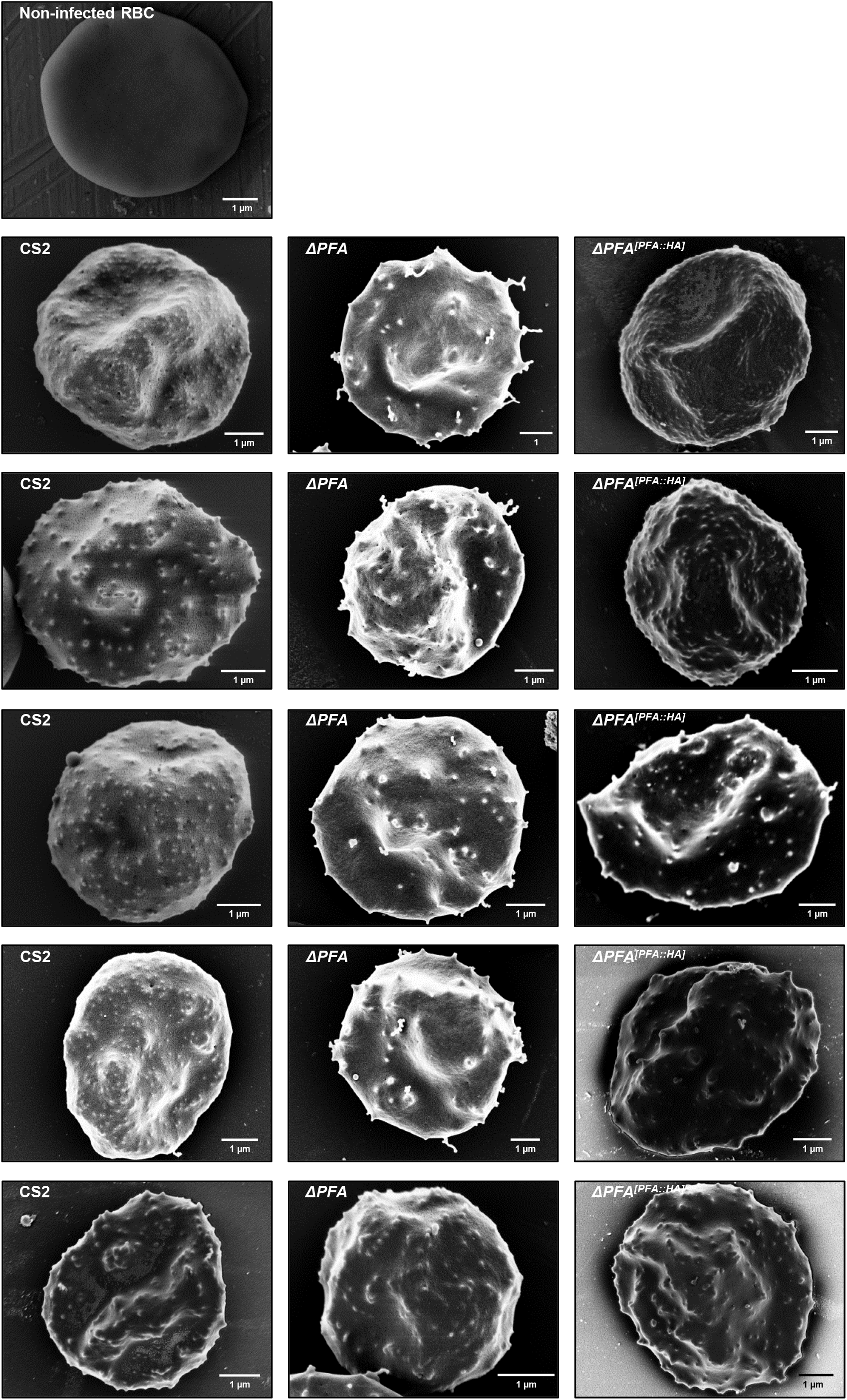
SEM image of a non-infected erythrocyte and additional SEM images of CS2, Δ*PFA* and Δ*PFA^[PFA::HA]^*.

**Figure S3.**
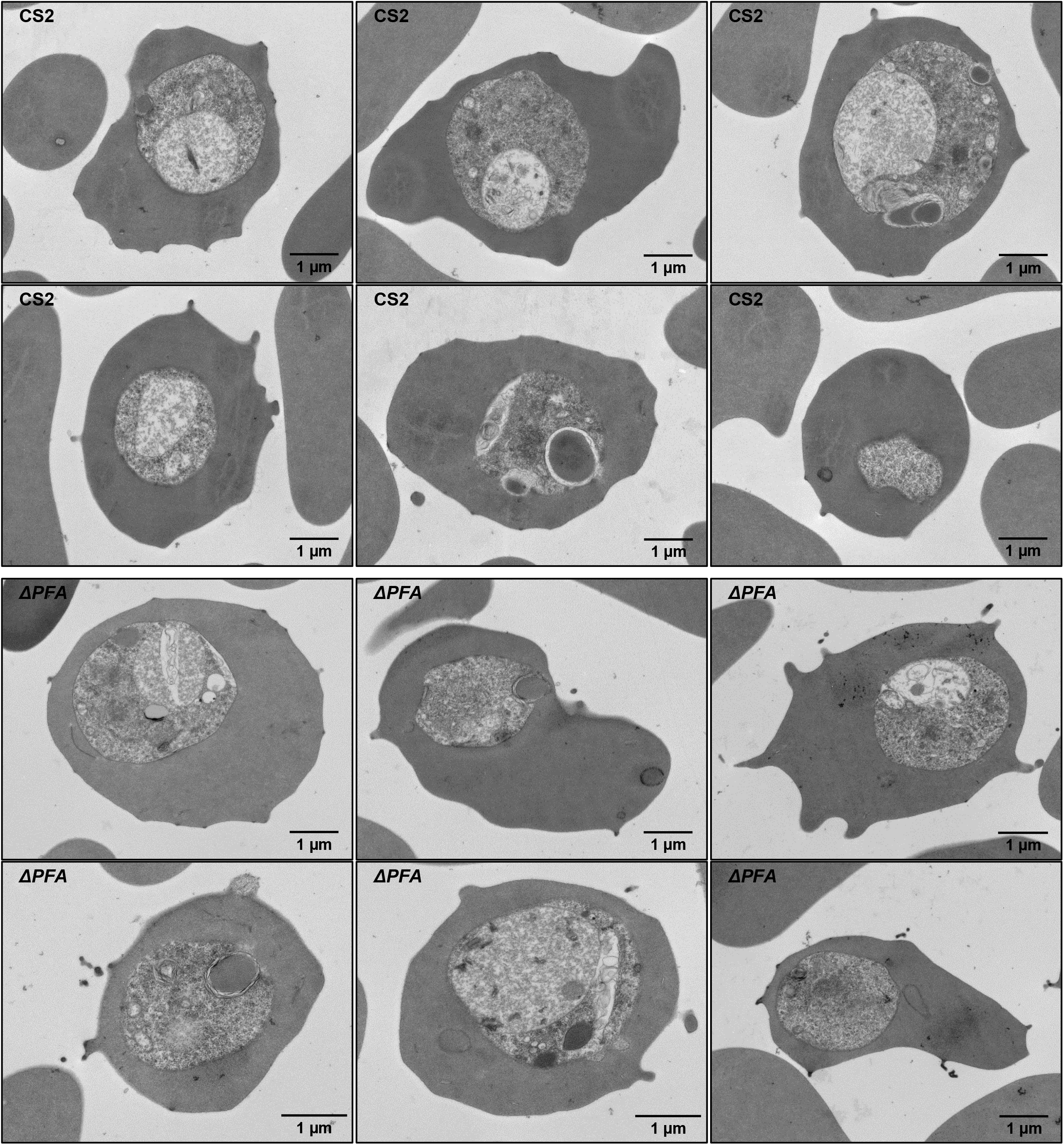
Additional TEM images of CS2 and Δ*PFA*.

**Figure S4.**
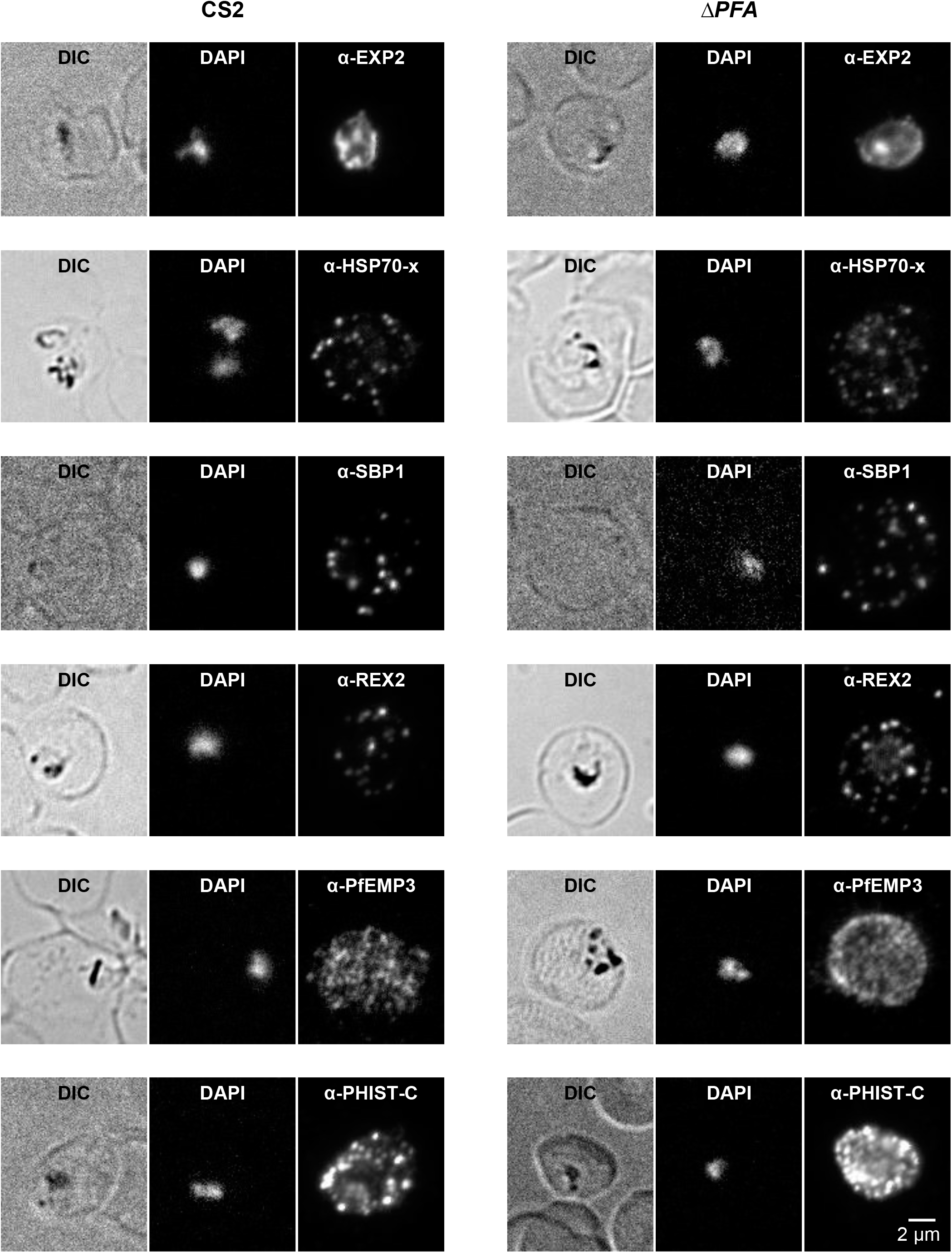
Investigation of marker protein localization using specific antisera in MeOH acetone-fixed Δ*PFA* with an immunofluorescence assay. No drastic difference in the localization of EXP2, HSP70x, SBP1, REX2, PFEMP3, or PHIST C was found.

**Figure S5.**
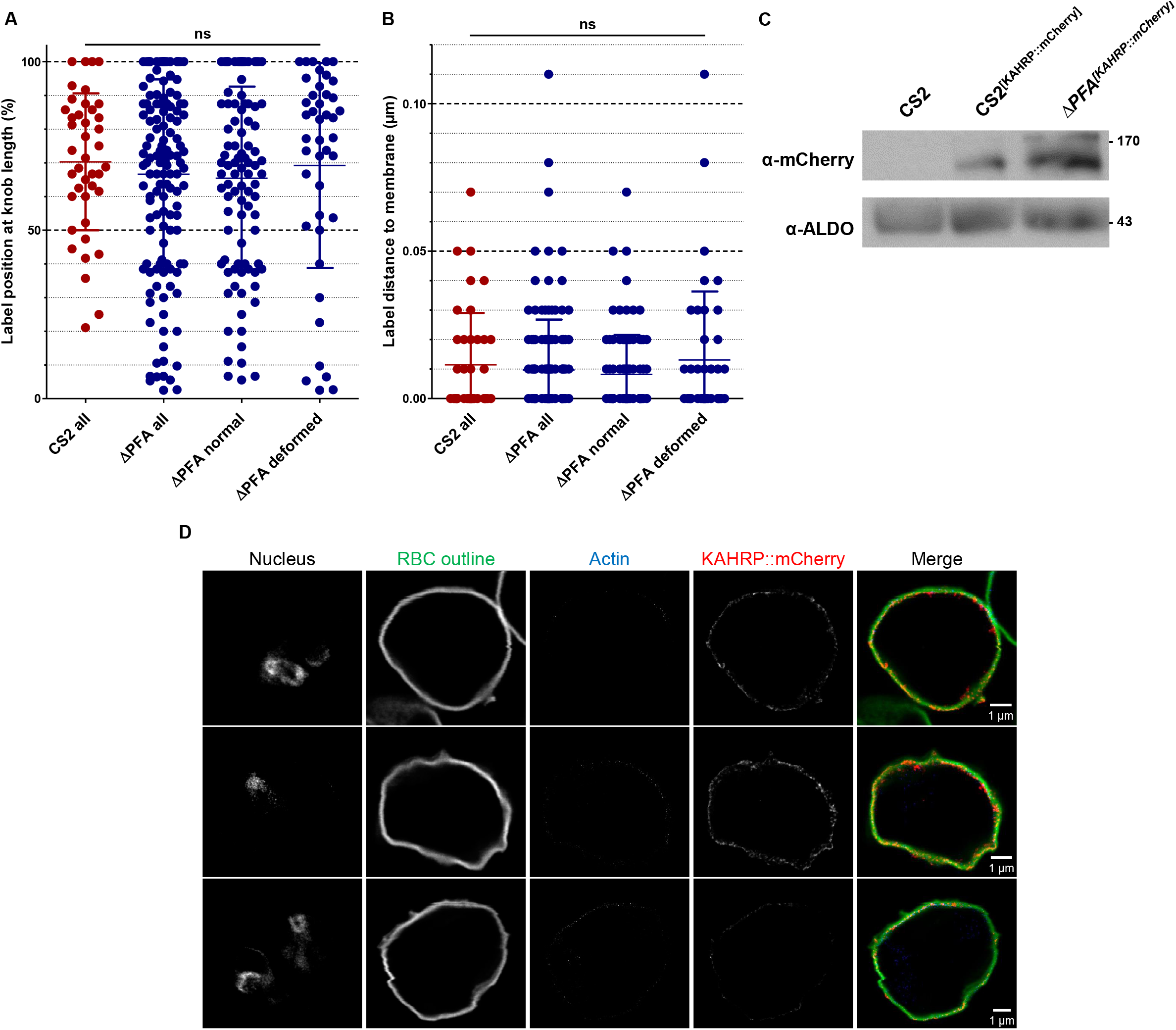
A) Investigation of label distribution in the α-KAHRP immuno-TEM. Distance of label from the base of the knob was measured using ImageJ and expressed relative to the length of the entire knob in percentages (0% being the base and 100% the top). The distribution of label along the full length of the knobs did not differ between the strains and knob types, B) Distance of label to the closest membrane was measured using ImageJ, revealing no difference between the strains and knob types. C) Verification of *CS2^[KAHRP::mCherry]^* and Δ*PFA^[KAHRP::mCherry]^* via Western blot verifies the production of KAHRP::mCherry protein in the cell lines using an α-mCherry antibody. The parasite protein aldolase (ALDO) was used as a loading control. D) RSTED images of *CS2^[KAHRP::mCherry]^* reveal close association of KAHRP::mCherry with the cytoskeleton and glycocalyx. Larger aggregates of KAHRP::mCherry were, in contrast to Δ*PFA^[KAHRP::mCherry]^*, not observed.

**Figure S6.**
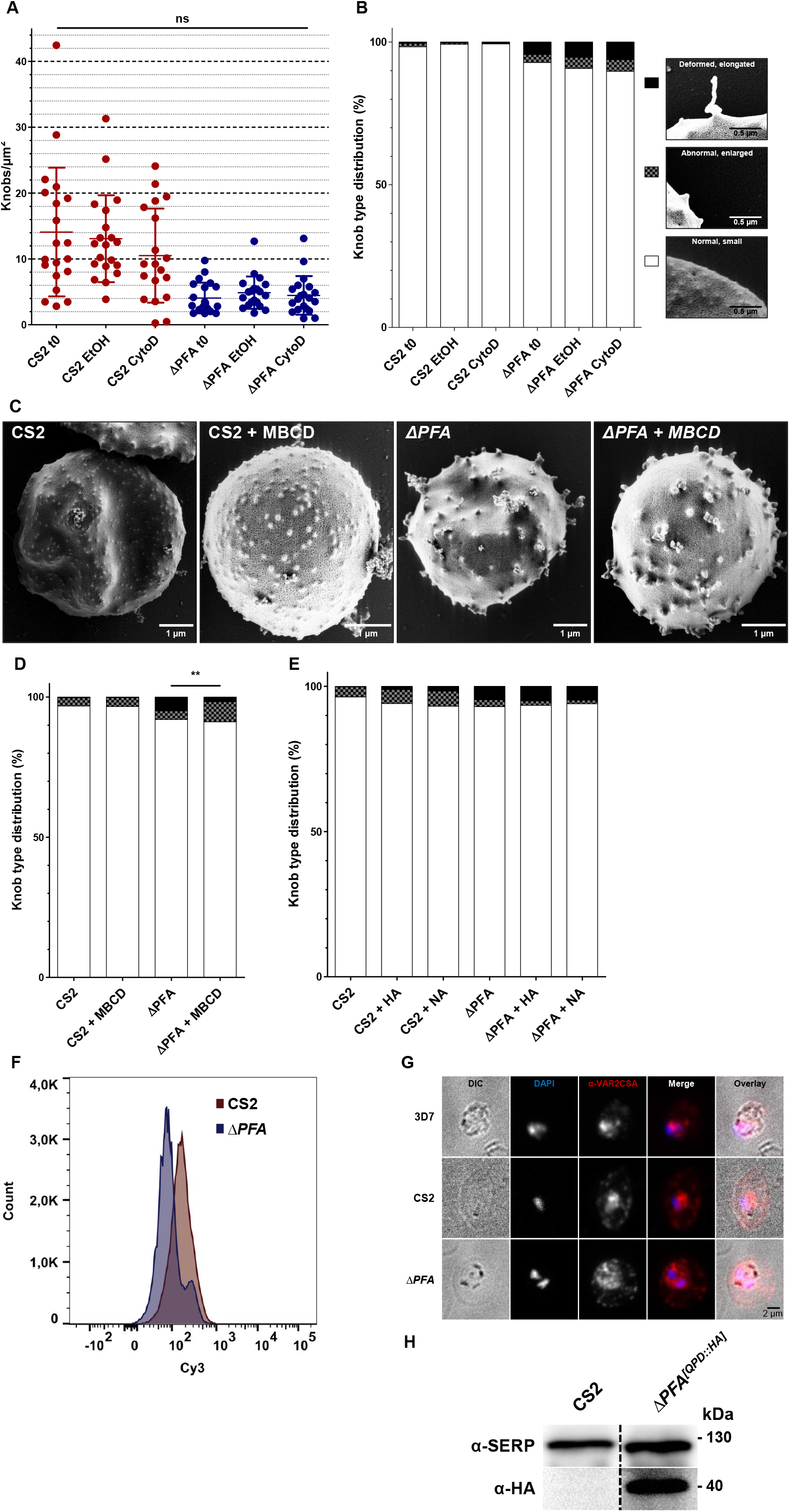
A) Treatment with the actin depolymerizing agent cytochalasin-D does not resolve knob density (A) and *mentula* knob morphology (B) N = 15. C, D) Investigation of treatment with the lipid raft disruptor MBCD (C, D) the glucosidases hyaluronidase (HA) and neuraminidase (NA) (E) on knob-type distribution in the two cell lines. N = 15. F) Concatenation of all iRBC data from the experiments in Figure 5B also shows a decrease of Cy3-caused fluorescence in Δ*PFA* across all experiments. Total number of single cells: 1,798,268 (CS2) and 1,799,274 (Δ*PFA*). G) Investigation whether MeOH-fixed parasites with an α-VAR2CSA antibody demonstrate that both CS2 and Δ*PFA* express var2CSA to similar levels. H) Western blot showing expression of QPD::HA in Δ*PFA^[QPD::HA]^* with an α-HA antibody. The parasite protein SERP was used as a loading control.

**Supplementary Video 1.** 3D reconstruction and surface render of *mentulae* depicted in Figures 2E, F. Red, erythrocyte plasma membrane; blue, electron dense material.

**Supplementary Video 2A.** Z-stack of erythrocytes infected with CS2^[KAHRP::mCherry]^ in mCherry channel.

**Supplementary Video 2B.** Z-stack of erythrocytes infected with CS2^[KAHRP::mCherry]^ in mCherry channel.

**Supplementary Video 2C.** Z-stack of erythrocytes infected with Δ*PFA*^[KAHRP::mCherry]^ in mCherry channel.

**Supplementary Video 2D.** Z-stack of erythrocytes infected with Δ*PFA*^[KAHRP::mCherry]^ in mCherry channel.

