## Supplementary material for "Co-chaperone involvement in knob biogenesis implicates host-derived chaperones in malaria virulence": Sup. Table 2A: Supplementary Table 2A.pdf

| Primer | Sequence (restriction sites underlined) |
|---|---|
| PFA_NotI_F | CAGCGGCCGCAACCTTAAGGAAAAGCTATG |
| PFA_MluI_R | ATACGCGTAGTAAATATATTATGATTTCCAGCAC |
| GFP_54_R | GTGCCCATTAACATCACCATC |
| Not-70_F | GGCGGATAACAATTTCACACAGG |
| KAHRP_F | GGCTCGAGATGAAAAGTTTTAAGAACAAAAATACTT |
| KAHRP_R | GGCCTAGGACCACAGCATCCTCTTTTCTTC |
| PFA0660w_5´_F | GACTTATTGAACTGCGGTTATATTATAAAG |
| PFA0660w_3´_R | GTATGCAAATCATAATAAATTCATATATC |
| HA_F | TAC CCG TAC GAC GTC |
| PFA_QPD_Quick_F | GCCATGAAATGGCAGCCTGATAAACATG |
| PFA_QPD_Quick_R | CATGTTTATCAGGCTGCCATTTCATGGC |
| mCherry_AvrII_F | GGCCTAGGATGGTGAGCAAGGGCGAGGAG |
| mCherry_XmaI_R | GATCCCGGGTTACTTGTACAGCTCGTCCAT |
